# Balance between resource supply and demand determines nutrient limitation of primary productivity in the ocean

**DOI:** 10.1101/064543

**Authors:** George I. Hagstrom, Simon A. Levin, Adam C. Martiny

## Abstract

What is the ultimate limiting nutrient in the ocean? The dominant theory, which was first proposed by Redfield and later formalized by Tyrrell[1, 2], states that despite the scarcity of fixed nitrogen (N) in the surface ocean, phosphorus (P) availability ultimately determines primary productivity. Two recent findings directly challenge the assumptions of the Redfield-Tyrrell paradigm: the discovery of systematic variations of phytoplankton cellular N:P:Fe and widespread iron-limitation of phytoplankton. Here we use a simple model of nitrogen, phosphorus, and iron (Fe) cycling to show how the resource demand ratios and biogeography of phytoplankton interact with external resource supply ratios to govern nutrient cycling and primary productivity. We find that all three nutrients can limit global primary productivity, and that the ratio of geochemical supply to biological demand of each nutrient in each ocean region determines the limiting nutrients, with nitrogen N fixation providing a mechanism for the cycles to interact. These results have important consequences for our understanding of biogeochemical cycles, ocean-atmosphere interactions, marine ecology, and the response of ocean ecosystems to climate change. Our work demonstrates the importance of resource ratios and suggests that future studies of the physiological and geochemical regulation of these ratios are indispensable to building accurate theories and future predictions of nutrient cycling and primary productivity.

---

The modern view of marine biogeochemistry is that N is the *proximate* and P is the *ultimate limiting nutrient* in the ocean. Initially, biologists used the results of bottle enrichment experiments as evidence that fixed N limits primary productivity, as populations showed dramatic responses to N but rarely to other nutrients[3, 4]. A much different line of reasoning came from geochemists, who pointed out that the marine fixed N inventory is regulated by N-fixation and denitrification, and that the availability of P may control the balance of these processes over geologic time-scales. These two ideas were reconciled by Tyrrell[2, 5, 7], who provided a simple mathematical model explaining how P-limited N-fixers can homeostatically regulate fixed N and primary productivity. This model explained the Redfield-Tyrrell paradigm[1] by suggesting that the deficit between the N:P of the deep ocean ((N:P)_d_) and organic matter ((N:P)_org_) is due to a N-fixer growth penalty, leaving other phytoplankton perpetually N limited. According to this viewpoint, N is the *proximate limiting nutrient* and P is the *ultimate limiting nutrient*.

Three observations challenge the Redfield-Tyrrell paradigm: widespread Fe-limitation of phytoplankton[8, 9], large scale systematic deviations of phytoplankton elemental stoichiometry[11, 12, 13], and restricted biogeography of N-fixers[29]. There is extensive evidence (as acknowledged by Tyrrell) that phytoplankton, particularly N-fixers, can be limited by Fe in large parts of the oceans. Furthermore, surface P levels in subtropical gyres are anti-correlated with estimates of aeolian dust flux and local dissolved Fe levels[14, 9], which suggests that N-fixers are only P-limited in high-Fe regions such as the subtropical North Atlantic Ocean[8]. Indeed Fe limitation of N-fixers is commonly observed throughout the oceans[20, 9]. Iron limitation of N-fixers points at Fe, and not P, as the *ultimate limiting nutrient*.

Latitudinal gradients in phytoplankton elemental composition challenge homeostatic models of ocean nutrient regulation. Recent surveys of (N:P)_org_[12] and nutrient tracers[13] show that (N:P)_org_ has systematic variations throughout the ocean: with (N:P)_org_ > 20 in subtropical gyres and (N:P)_org_ < 12 in high-latitude regions and highly productive systems. These patterns are not yet fully understood[6], but appear to be set by some combination of nutrient levels[15], phytoplankton composition[12], and/or temperature[16]. Differences in elemental stoichiometry between different species of phytoplankton may also be important: N-fixers have N:P and Fe:P greater than non-fixers[19], suggesting that whether diazotrophs are limited by Fe or P may alter their interaction with regular phytoplankton. Furthermore, N-fixers are geographically restricted to low-latitude, usually oligotrophic ocean regions in which N:P ratios are significantly above the Redfield ratio. As a result, we expect that changes in phytoplankton nutrient demand would impact the role of N-fixers and more broadly patterns of nutrient limitation.

## Box-Model Framework

Building on the framework outlined by Tyrrell, we constructed a simple box model with only 11 variables, incorporating the cycling of three nutrients, three phytoplankton types, and regional differences in ecosystem structure, to illustrate the importance of both biogeography, Fe-cycling, and nutrient supply and demand ratios. The model structure and is shown in Figure 1.

**Figure. 1.**
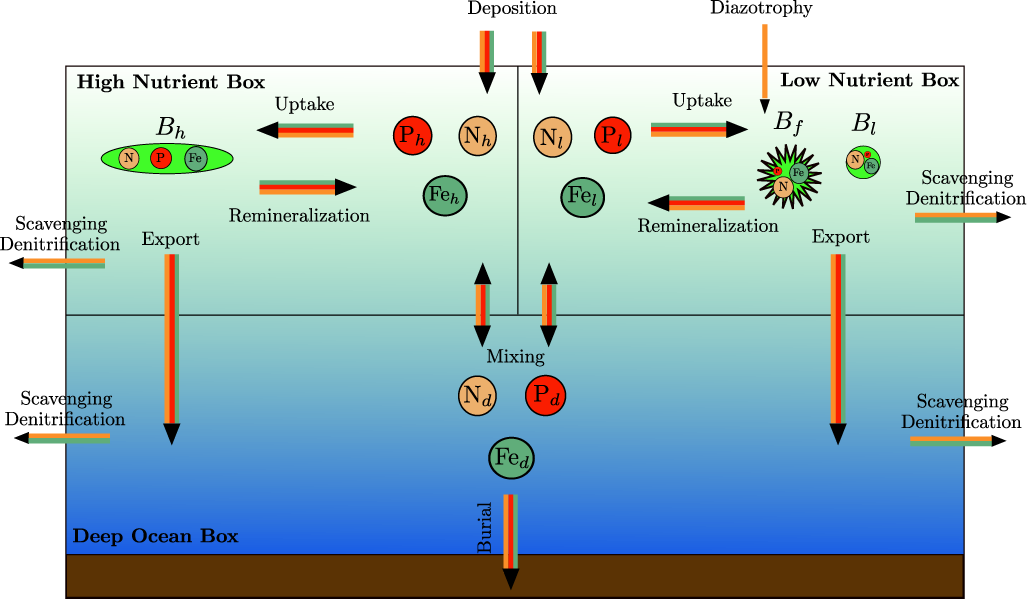
Diagram of biogeochemical model of N, P, and Fe cycles. Variables in the high-nutrient supply box (denoted with *h*) represent nutrients and biomass in ocean regions with high levels of mixing, such as equatorial upwelling systems or subpolar gyres. Variables in the low-nutrient supply box (denoted with *l*) represents nutrients and biomass in subtropical gyres. Three phytoplankton types are modelled-two non-fixers living in the high- and low-nutrient regions, and a N-fixer living in the low-nutrient supply region.

The model includes two surface boxes to capture ecological and physical differences between regions with high-nutrient supply, such as sub-polar gyres and equatorial or coastal regions, and low-nutrient subtropical gyres. We model the differences in nutrient supply and phytoplankton stoichiometry between the boxes, and restrict N-fixers to the low nutrient-supply box, which is consistent with their biogeography [29]. We cover the dominant processes influencing the nutrient availability of each element, including the loss of P in the deep ocean, Fe-scavenging and complexation, and denitrification. Our model resolves nutrient dynamics on time scales up to 10^5^ years, which is roughly the residence time of P in the ocean. Because it is a box model, it cannot generate precise predictions about ocean dynamics, but instead provides a low-dimensional description of biogeochemical cycles, identifying the most important mechanisms governing productivity in the ocean.

## Results

On the short time-scale of surface mixing, deep nutrient concentrations remain approximately constant. In contrast surface nutrient concentrations and phytoplankton biomass are dynamic and rapidly converge to *proximate* equilibrium states determined by the rate of nutrient supply from deep ocean and atmospheric sources. On much longer time-scales, deep ocean N and P (N_*d*_ and P_*d*_) change, leading to slow shifts in nutrient supply and in the surface equilibrium states. We used these links between nutrient supply, productivity, and N-fixation to to determine how external nutrient fluxes control the biogeochemical equilibrium of the N and P cycles, to determine the ultimate limiting nutrient(s), and the controls on deep ocean resource ratios.

We first consider the short time-scale dynamics in the low-nutrient box with N_*d*_ and P_*d*_ held constant. This leads to chemostat equations for N-fixers and non-fixers competing to grow on N, P, and Fe. N, P, and Fe are supplied at the constant rates 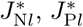, and 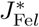, which are determined by mixing and external deposition, so that 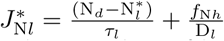, with analogous expressions for P and Fe flux. The starred concentrations are the *R\** values of high-nutrient phytoplankton derived from resource competition theory[31], *τ*_*l*_ is the low-nutrient mixing time, and *f*_N*l*_ is the external supply of N to the low-nutrient box. Due to a growth penalty, N-fixers have higher values of Fe* and P* than non-fixers, so that coexistence between the two types at equilibrium is only possible when non-fixers are limited by N. Therefore, we determine the *proximate limiting nutrient* (PLN), whose concentration determines non-N-fixer growth rate, by comparing the supply of N, P, and Fe with non-fixer stoichiometry:

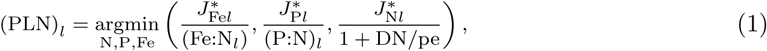

Here pe is the fraction of primary productivity exported to the deep. If the PLN is N, then N-fixers may coexist with non-fixers, and can be limited by either P or Fe. How does low-nutrient primary productivity (TPP_*l*_) depend on nutrient supply? TPP_*l*_ is controlled by the supply rate of the nutrient that limits N-fixers and the average stoichiometry of the community with respect to that nutrient, which is determined by the ratio of N-fixer to non-fixer biomass. This is calculated from N mass-balance: each unit of Fe or P consumed by N-fixers contributes 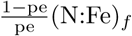 or 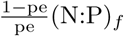 units of N for non-fixers. Nitrogen balance is achieved when non-fixer productivity equals the sum of available N from the atmosphere and ocean and N-fixery, so that in proximate equilibrium:

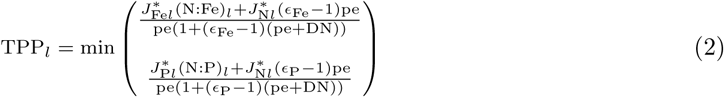

Here 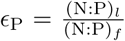 and 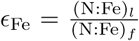 are the relative efficiencies of N-fixer P and Fe utilization. Equation 2 indicates that in surface equilibrium, both the stoichiometry of non-fixers and N-fixers determine the sensitivity of TPP_*l*_ to Fe- or P-flux from both the atmosphere and ocean mixing, and that TPP_*l*_ is a function of the N-flux as long as N-fixers and non-fixers have different stoichiometry. When diazotrophs and non-diazotrophs have the same stoichiometry, this result suggests that the effect of a sudden increase in N-flux is damped much faster than indicated in other studies[2], on the time-scale of surface equilibrium rather than the the equilibrium of the N-cycle.

Figure 2 shows how N-fixer stoichiometry influences the sensitivity of TPP_*l*_ to nutrients. Because N-fixer Fe-use is inefficient, 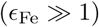[19], increases in N-supply shift the community to higher N:Fe and TPP_*l*_. When they are P-limited, TPP_*l*_ is insensitive to N-supply since N-fixers use P as efficiently as non-fixers. N-fixers are limited by the nutrient that minimizes Equation 2, which is a function of Fe and P supply and demand.

**Figure. 2.**
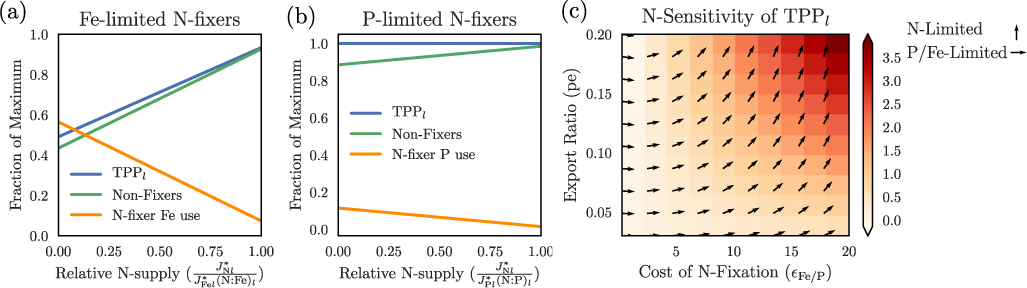
Relative influence of N and Fe supply on TPP_*l*_ under proximate limitation. Panels (a) and (b) show how increases in 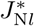 change TPP_*l*_, non-fixer biomass (B_*l*_), and N-fixer use of Fe (a) or P (b). Panel (c) shows the sensitivity of TPP_*l*_ to N and P/Fe, in terms of pe and the cost of N-fixery ∊Fe or ∊P. The angle of each arrow is that of a vector with components proportional the gradient of TPP*l* with respect to 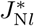 and 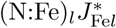, so that arrows indicate sensitivity to N only, rightward arrows indicate sensitivity to P/Fe only.

Next, we find that total primary productivity in the high-nutrient box is given directly by the supply rate of the nutrient that is most scarce compared to the stoichiometry of high-nutrient box phytoplankton:

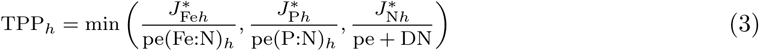

Here the proximate limiting nutrient in the high-nutrient box is the nutrient which minimizes Equation 3.

Finding the *ultimate limiting nutrient* (ULN) is a two-step process: given the external fluxes of each nutrient to each box, we first determine which nutrient limits each phytoplankton type. Through the formulas for TPP in each box in the proximate-equilibrium state, we know the dependence of TPP on the rates of nutrient supply (i.e. 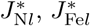, etc). A nutrient is an *ultimate limiting nutrient* if its external supply rate (i.e. *f*_N*l*_, *f*_Fe*l*_, etc) influences TPP at biogeochemical equilibrium, which we diagnose by studying the sensitivity of the nutrient supply rates to the external nutrient fluxes.

Fe and P supply rates depend simply on the external fluxes, whereas the N supply can depend on external fluxes of all three nutrients. Fe has a short residence time in the ocean and the supply of Fe to each ocean region is sensitive only to the external Fe flux to that region (*f*_Fe*h*_ and *f*_Fe*l*_). At biogeochemical equilibrium, P_*d*_ is nearly equally sensitive to external P flux from each region, and is the quotient of the total external P supply and the product of ocean volume and the P loss rate constant. Because P supply rates are dominated by the supply from the deep ocean, they are equally sensitive to low- and high-nutrient external fluxes (*f*_P*l*_ and *f*_P*h*_).

The rate of N supply to each ocean region depends strongly on the concentration of deep ocean N at biogeochemical equilibrium. Equilibrium in the N-cycle is reached when gains from N-fixery and external fluxes match losses from denitrification. The rate of N-fixation depends on the biomass of diazotrophs, and is therefore a function of the supply rate of both N and either Fe or P to the low-nutrient box. Thus, N-supply rates can be influenced by any of the nutrient fluxes, depending on which nutrients limit N-fixers and high-nutrient phytoplankton.

Our analysis illustrates that any of the three nutrients can be the *ultimate limiting nutrient*. TPP has a simple relationship to the supply rate of the nutrients limiting all three types of phytoplankton, which allows us to determine the ULNs. The analytical expressions for TPP in each box (Equations 2 and 3) directly indicate that P or Fe can be an ULN, if either limits N-fixers or high-nutrient phytoplankton. If diazotrophs are Fe limited, then TPP is strongly dependent on the low-nutrient Fe flux (*f*_Fe*l*_) and Fe is an ULN. If they are P limited then TPP is strongly dependent on both P fluxes, and P is an ULN. A similar picture holds within the high-nutrient box: high-nutrient Fe flux strongly influences TPP when the high-nutrient phytoplankton are Fe limited, and both P fluxes increase TPP when high-nutrient plankton are P-limited.

Figures 3 (a) and (b) show how nutrient supply and demand control nutrient limitation patterns by examining the effects of Fe deposition on TPP. When external flux of Fe to the low-nutrient box (*f*_Fe*l*_) is weak, N-fixers are Fe-limited and N_*d*_ is low. Iron will be an ULN in the low-nutrient region, and N will limit phytoplankton in the high-nutrient region. As low-nutrient box Fe deposition is increased, N_*d*_ increases and there is an N-teleconnection between the low and high-nutrient regions, leading to a transition from N to Fe-limitation of the high-nutrient regions, so that Fe and potentially P are the ULNs. Further increases in low-nutrient Fe lead to a transition to P-limitation in the low-nutrient box. As high-nutrient box Fe supply is increased, there is a transition from Fe to N- or P-limitation in the high-nutrient region. The dashed contours in the figure correspond to places in parameter space where the relative supply and demand of two different nutrients are equal, indicating potential transitions from different nutrient limitation scenarios.

**Figure. 3.**
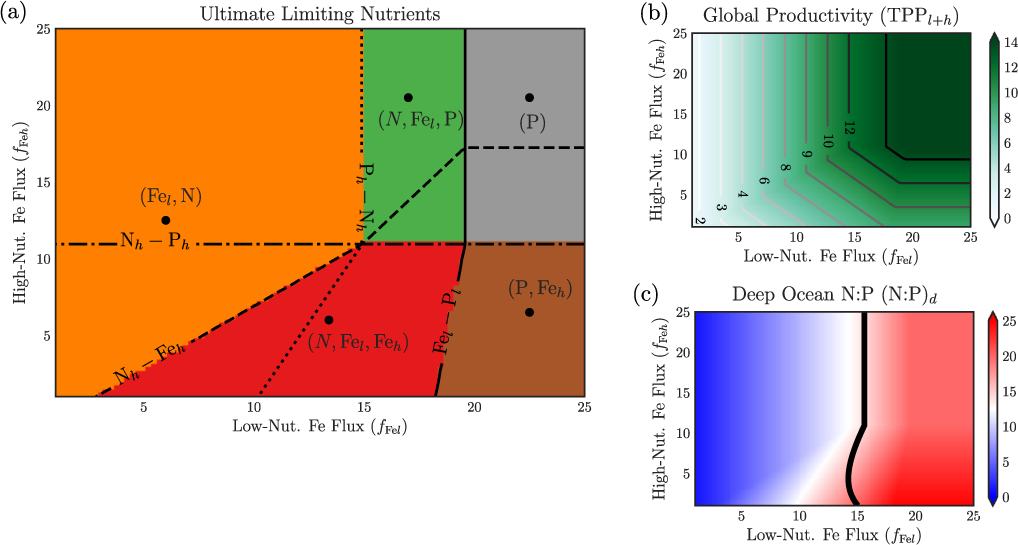
Influence of varying ocean conditions on nutrient limitation (a) and primary productivity (b) and stoichiometry (c). Figure (a) shows the ultimate limiting nutrients as a function of Fe-deposition rates in both high- and low-nutrient boxes. The black contours show the points where relative supply and demand for two different nutrients are equal to eachother. Figure (b) shows the impact of Fe-levels on TPP. Figure (c) shows (N:P)_d_ as a function of Fe deposition. The black contour is the set of points where (N:P)_org_ = (N:P)_d_.

Can N be an ULN? The analytical expressions for TPP*_l_* and TPP_h_ (equations 2 and 3) demonstrate that TPP is a function of the rate of N-supply to both boxes, and that this dependence is particularly strong when N-fixery are both Fe-limited and require a large investment in Fe, and when high-nutrient box plankton are N-limited. But in order for N to be an ULN, the N-supply must also be strongly dependent on the external N-fluxes *f*_N*l*_ and *f*_N*h*_. We directly calculated the sensitivity of both low- and high-nutrient box TPP to external N-fluxes by taking the derivatives of TPP at bigeochemical equilibrium. The external N-flux to both boxes can have a significant impact on TPP. Ignoring less important parameters, we find that the external N-flux to both boxes can have a significant impact on TPP:

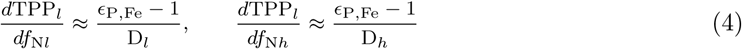

Here D*_l_* and D*_h_* are the depths of the low- and high-nutrient boxes.

The explicit dependence of TPP_*l*_ on atmospheric N deposition arises due to differences in N-fixer and non-fixer stoichiometry[18], introduced in the discussion of TPP_*l*_ on short-time scales. When N-fixers are Fe-limited, each unit of diazotroph productivity represents a lost opportunity for non-fixers to use Fe at a much higher N:Fe. Sustained increases in external N-fluxes shift Fe-utilization in the low-nutrient box, leading to lower N-fixer populations and higher TPP at equilibrium (Figure 4 (a)). This demonstrates the sensitivity of TPP_1_ to external Fe and N. When N-fixers have a high Fe content, the external N-flux has a significant impact on primary productivity, and N becomes an ultimate limiting nutrient. This effect vanishes only in the unlikely scenario that N-fixers and regular phytoplankton have the same stoichiometry (∊ = 1), in which model solutions are analogous to those found by Tyrrell.

**Figure. 4.**
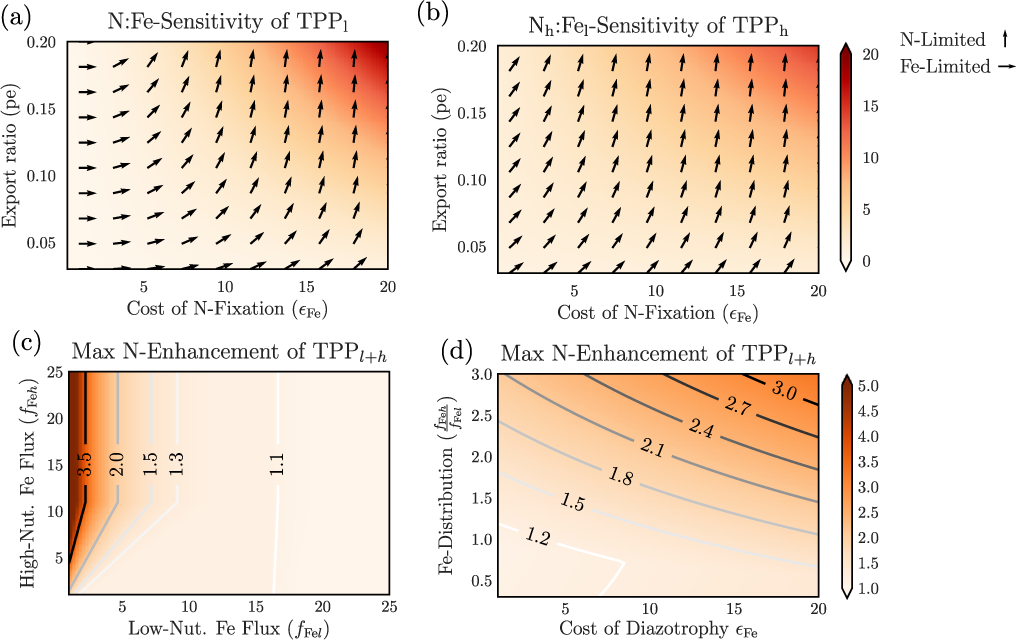
N-controls on TPP*_l_* and TPP*_h_* at biogeochemical equilibrium. Panels (a) and (b) show the sensitivity of TPP*_l_* and TPP*_h_* to external N and Fe deposition, normalized to (N:Fe)*_l_*, with respect to the pe-ratio and ∊Fe, when N-fixers and high-nutrient plankton are Fe- and N-limited, respectively. Panels (c) and (d) show the multiple by which global TPP would increase if N was replete, varying levels of atmospheric iron flux in (c), and the cost of N-fixery and distribution of iron between high and low nutrient boxes in (d).

The lack of N-fixers in high-nutrient regions decouples the N-cycle from other nutrient cycles here. High-nutrient phytoplankton are commonly N-limited[10], and our model suggests that when they are N-limited, TPP will be highly sensitive to N-fluxes, in particular the high-nutrient box flux:

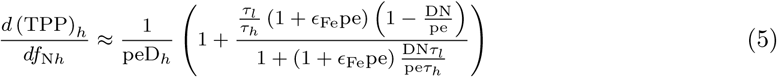

Equation 5 contains a term proportional to 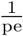, which makes TPP_*h*_ highly sensitive to high-nutrient box N-deposition (f_*Nh*_) no matter ∊Fe, as is seen in Figure 4 (b). When high-nutrient box phytoplankton are N-limited, N-additions can lead to several-fold increases in TPP at biogeochemical equilibrium. Our finding that N can be an ultimate limiting nutrient despite the presence of N-fixation is a significant departure from previous theoretical models, which were unable to capture the role of N due to assumed Redfield stoichiometry and lack of representation of N-fixer biogeography. Hence, we find that each nutrient has the potential to be an *ultimate limiting nutrient*, and most solutions lead to multiple nutrients simultaneously controlling TPP.

What happens to TPP when there are large, sudden changes in the nutrient fluxes? We used numerical experiments to study the dynamic response of the ocean to increases in Fe, N, and P fluxes. The model parameters of each experiment are marked by the circles in Figure 3 (a), where the red circle is representative of modern ocean conditions as determined by TPP and (N:P)_d_, and the initial condition corresponds to the stable equilibrium state with those parameters. The equilibrium state was perturbed by either doubling the Fe or P fluxes or by increasing N-flux to modern levels. As predicted by our theory, TPP strongly responds to sudden increases in the supply of the ultimate limiting nutrients, and more than one nutrient can be the ULN. Figures 5 (a) and (d) (which corresponds to the modern ocean) show dramatic increases in TPP under Fe addition, as was hypothesized by Martin[21] and Falkowski[8], as well as N addition, which is at odds with current thinking. Phosphorus additions can also stimulate TPP, as happens in Figures 5 (b), (c), and (e).

**Figure. 5.**
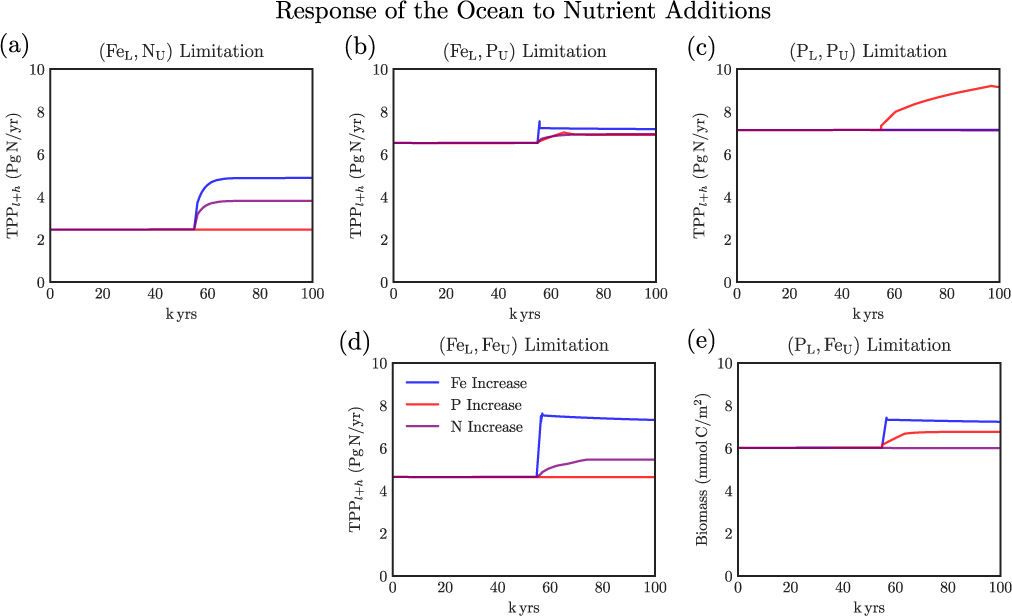
Response of the ocean to increases of the N, P, and Fe fluxes in different ultimate nutrient limitation regimes. Each figure corresponds to a numerical experiment with parameters selected from the marked locations in Figure 2. Figure (d) corresponds to both the (Fe, Fe) limitation regime and our best estimate of the modern ocean.

The primary geographic restriction of N-fixers to low-nutrient regions means that deep ocean nutrients are tied to the stoichiometry of phytoplankton in the low-nutrient regions. We demonstrated this by calculating the Fe-flux controls on deep ocean N inventory (Figure 3 (c)), and with explicit formulas linking deep-ocean N with the supply of each nutrient:

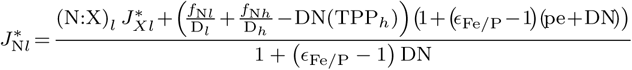

This equation shows the links between deep ocean N, the supply of whichever nutrient limits N-fixers, and the stoichiometry of N-fixers and non-fixers. There is homeostatic connection between the supply of N and either Fe or P, whose ratio is tied to the stoichiometry of low-nutrient phytoplankton. It further demonstrates that the strength of this homeostasis is modulated by the nutrient supply in the high-nutrient regions and thus to the ratio of high- and low-nutrient mixing times 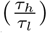. When 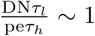 ∼ 1, deep ocean resource ratios cannot be maintained close to low-nutrient box phytoplankton stoichiometry. In the specific case of Fe-limited N-fixers, the gap between realized TPP*_l_* and the maximum TPP*_l_* sustainable by the low-nutrient Fe supply can be bridged by external N deposition (Figure 4 (c) and (d)), which shows the potential for external N-deposition to increase TPP several fold. Our results indicate that N_*d*_ is primarily controlled by either the N:P or N:Fe ratio of non-fixer in the low-nutrient box, so that the observation that deep ocean N:P is much below the N:P of organic matter in oligotrophic gyres suggests that the ultimate limiting nutrients are Fe and N and that deep ocean N:P ratios are not regulated by the mean N:P of phytoplankton.

## Discussion

Our study differs from prior theoretical investigations of the N, P, and Fe cycles. Tyrrell[2], Lenton[5, 7] and Augeres[33] modeled the N and P cycles using box models, assuming homogeneity of biological and geochemical variables in the surface ocean. These studies reproduced the Redfield-Tyrrell paradigm, finding that P is the ULN, and that (N:P)_d_ is homeostatically controlled by (N:P)_org_. Our results diverge from these earlier studies because we modeled surface ocean heterogeneity, which allowed us to capture the covariance between nutrient concentrations, phytoplankton stoichiometry, N-fixer biogeography, and nutrient supply from both ocean mixing and external deposition. Box models rely on averaging of higher-resolution equations over the boxes and deriving approximate equations for the means of the dynamical variables in each box. Our selection of boxes isolates ocean regions by their biological and geochemical properties, minimizing-within box covariances and enabling a much more accurate representation than prior models. This seemingly small change led to the large differences in our understanding of ocean biogeochemistry.

Weber and Deutsch[32] found that *circulation averaging* could maintain homeostasis between (N:P)_d_ and (N:P)_org_ if there is significant lateral transport of P-deplete waters from high-into low-nutrient regions, (see also SI section S5). For circulation averaging to maintain this homeostasis, the N:P in laterally transported waters must be significantly above the Redfield ratio. However, direct observations[30] suggest that the N:P of nutrients transported laterally into oligotrophic gyres is much below (N:P)_org_. This elevated N:P is likely due to efficient remineralization of P compared to N. If Fe is replete, both lateral transport of low N:P nutrients and efficient P remineralization would drive (N:P)_d_ much higher than the (N:P)_*l*_ (SI section S3 and S4), strongly enhancing the outcome where Fe and N are ULNs.

Our simplified model of marine biogeochemical cycling revealed the most important controls on biogeochemical cycling at the cost of quantitative accuracy. Box models are limited because they grossly simplify biological and physical ocean dynamics, so we do not expect them to make quantitatively accurate predictions. Nevertheless, the structure of our box-model was chosen carefully to minimize the covariances of biological and geochemical properties within each box, and therefore we expect that our model identifies the most important factors influencing nutrient inventories and primary production. Our model treats the nutrient ratios of different phytoplankton types as static features. However, marine ecosystems are complex adaptive systems[34] and phytoplankton nutrient demands are thus subject to change by physiological acclimation[24] or ecological/evolutionary selection[25, 15]. Our parametrization of phytoplankton stoichiometry closely mirrors field and laboratory observations[6], and we should expect a more realistic model to have similar properties[33], but there may be some effects that cannot be captured by a purely static model. For example, phytoplankton may be frugal[26] in their resource utilization, increasing or decreasing their quotas in response to availability. Such co-evolution of supply and demand would drive the ocean towards colimitation by all three nutrients by shifting both non-fixer and N-fixer resource ratios to enhance utilization of any excess nutrients.

We found that external N-fluxes can play a major role in influencing TPP even on biogeochemical time-scales[36], which was thought to be impossible when there are strong feedbacks controlling N-fixation[35], such as under the Redfield-Tyrrell paradigm. This could occur in two ways: through N-limitation of phytoplankton in high-nutrient regions, or through Fe-limitation of N-fixers, so that the influence of N-fluxes depends strongly on phytoplankton resource ratios and nutrient limitation patterns. Anthropogenic inputs of N have become highly significant since industrialization[22], and these fluxes may already be comparable to N-fluxes from N-fixers, leading to large-scale eutrophication of coastal regions and shallow seas. Our work suggests that these inputs may have a much more widespread impact, and that they have the potential to alter primary productivity on a global scale. In order to fully understand the role of anthropogenic-N in marine ecosystems, we require a deeper understanding of the patterns of phytoplankton nutrient limitation and the factors controlling resource ratios of N-fixers and phytoplankton.

## Materials and Methods

### Ecological Model

The concentration of N-fixers, non-fixers, and large phytoplankton are the variables *B_f_*, *B_p_*, and *B_h_*. N-fixers and non-fixers are restricted to the low-nutrient box, and large phytoplankton are restricted to the high-nutrient box. Each type is modeled with the Monod equation and Liebig’s law of the minimum[31]. A type is specified by its maximum growth rate, half saturation constants, and nutrient stoichiometry. Mortality rates due to all processes are assumed to be constant, and dead phytoplankton are instantly remineralized into organic nutrients with a small fraction exported into the deep-ocean. These variables (measured in mol N) evolve according to:

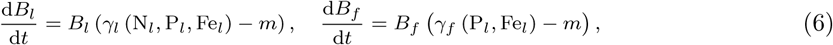

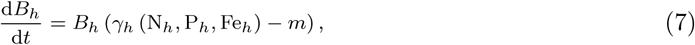

Here the growth rate terms are:

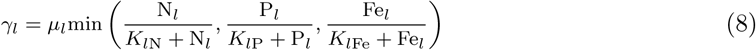

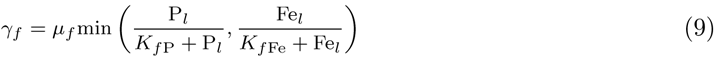

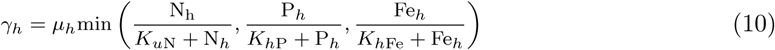

The N-fixer growth rate *γ*_*f*_ does not depend on *N_l_* in order to reflect the fact that N-fixers utilize dissolved N_2_ instead of fixed N. The explicit representation of the N:P:Fe ratios of each phytoplankton type allow us to determine the sensitivity of important biogeochemical processes to phytoplankton stoichiometry. Here we focus on an ecologically realistic parameter range, (N:P)_*f*_ > (N:P)_*l*_ > (N:P)_*h*_ and (Fe:N)_*f*_ > (Fe:N)_*h*_ ∼ (Fe:N)_*l*_.

### Phosphorus Cycling

The inorganic P concentrations in the low-nutrient, high-nutrient, and deep-ocean regions are P_*l*_, P_*h*_, and P_*d*_. The concentration of P is affected by ocean transport, nutrient uptake and remineralization, external deposition, and burial. Burial is modeled as a first-order loss process from the deep-ocean. P uptake is determined by the rate of phytoplankton growth and phytoplankton stoichiometry. Remineralization leads to a flux of nutrients into the surface and deep-ocean boxes proportional to the mortality rate, the total biomass, the elemental stoichiometry of each phytoplankton type, and the rates of remineralization. Transport between boxes is modeled by means of exchange fluxes, as is typical in box-modeling studies. The equations for P are:

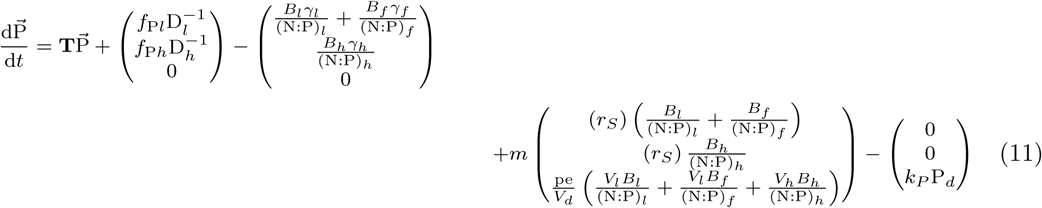

where the transport matrix **T** and the P vector 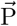 are:

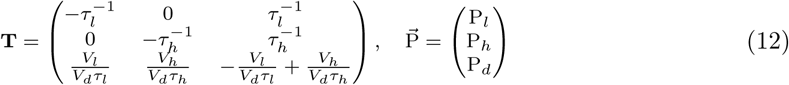

### Nitrogen Cycling

The total concentration of fixed inorganic N in the high-nutrient, low-nutrient, and deep-ocean boxes are represented by the variables N_*l*_, N_*h*_, and N_*d*_. New N is supplied by external deposition and N-fixation. Denitrification is modeled as a loss of a fixed percentage of N during remineralization. We exclude sea-floor burial terms because N fixation and denitrification are the dominant processes. The equations for the N concentrations in each box are:

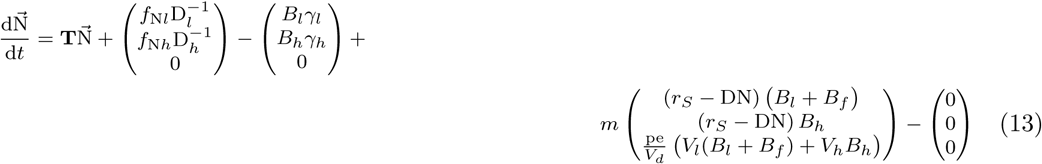

### Iron Cycling

Dissolved Fe(2) is rapidly oxidized to Fe(3) and lost via scavenging. Organic ligands form complexes with Fe(2) and prevent scavenging, and the majority of dissolved Fe exists in complexed form. Bioavailable Fe is the sum of dissolved Fe and complexed Fe (Fe*), represented by Fe_*l*_, Fe_*h*_, and Fe_*d*_. Scavenging of Fe is modeled following Parekh[27]: there is assumed to be a constant global concentration of organic ligands, *L_T_* = 10^−9^ mol/L, and the equilibrium constant for complexation with dissolved Fe is κ_*Fe*_ = 10^11^. The equilibration of dissolved/complexed Fe and free ligand is treated as instantaneous: 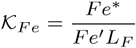, where Fe′ is dissolved Fe and L_*F*_ is the free ligand. Scavenging is a loss term proportional to the uncomplexed dissolved Fe, and is proportional to *J*(Fe) = *J_sc_*Fe′(Fe). We fix the deep-ocean Fe concentration to be Fe_*d*_ = 5 × 10^−10^mol/L, which is justified by deep ocean ligand concentrations. As a result, there are two equations for the time evolution of the surface Fe concentrations, which include contributions from external deposition:

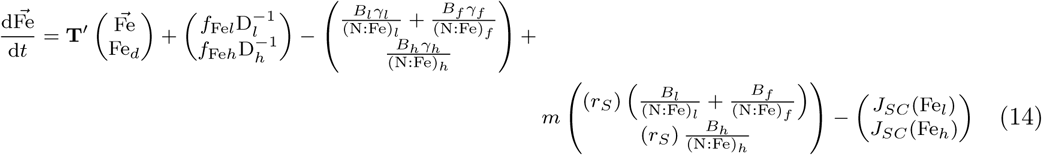

Here we have defined the reduced transport matrix:

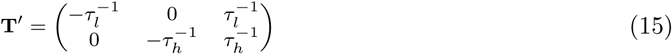

## Acknowledgements

We acknowledge Francois Primeau for providing thoughtful discussion as well as suggestions. This work was partially supported by NSF Dimensions of Biodiversity grant OCE-1046001 and OCE-1046297 (S.A.L., A.C.M.), Army Research Office Grants W911NG-11-1-0385 and W911NG-NF-14-1-0431 (GIH, SAL), and Simons Foundation Grant 395890. The authors do not have any competing financial interests.

